# A mosquito parasite is locally adapted to its host but not temperature

**DOI:** 10.1101/2023.04.21.537840

**Authors:** Kelsey Lyberger, Johannah Farner, Lisa Couper, Erin A. Mordecai

## Abstract

Climate change will alter interactions between parasites and their hosts. Warming may affect patterns of local adaptation, shifting the environment to favor the parasite or host and thus changing the prevalence of disease. We assessed local adaptation in the facultative ciliate parasite *Lambornella clarki*, which infects the western tree hole mosquito *Aedes sierrensis*. We conducted laboratory infection experiments with mosquito larvae and parasites collected from across a climate gradient, pairing sympatric or allopatric populations across three temperatures that were either matched or mismatched to the source environment. *L. clarki* parasites were locally adapted to their hosts, with 2.6x higher infection rates on sympatric compared to allopatric populations, but were not locally adapted to temperature. Infection peaked at the intermediate temperature of 13°C. Our results highlight the importance of host selective pressure on parasites, despite the impact of temperature on infection success.

## Introduction

Climate change is likely to have a significant impact on species interactions such as those between hosts and parasites. In particular, warming temperatures have been associated with changes in the prevalence and intensity of disease outbreaks (Altizer et al 2013, Lafferty and Mordecai 2016). Variation in the thermal sensitivities of parasite populations may play an important role in determining where warming will have the largest effect. Specifically, parasite populations are often locally adapted, performing best in climates similar to their home environments. For example, some populations of chytrid fungus show a signal of local adaptation to temperature (Stevenson and Pike 2013). Species exhibiting local thermal adaptation – that is, standing variation in thermal tolerance that corresponds to the source environment – may be better able to adapt to warming and persist under climate change. Conversely, species with minimal between-population variation in thermal tolerance may have a reduced capacity to adapt. Understanding not only the sensitivities of parasites and pathogens to temperature, but also the variation in thermal sensitivities among populations, is critical for predicting how climate change will impact disease dynamics (Sternberg and Thomas 2014).

Parasites and pathogens may also be locally adapted to their hosts. Hosts and parasites are engaged in a reciprocal evolutionary interaction (i.e, an “evolutionary arms race”) in which hosts evolve immune or behavioral defenses and parasites must evolve to overcome them. The outcome of host – parasite coevolution, whether parasites or hosts are winning the race, depends on generation time, genetic variation, and dispersal (Gandon 2002, Gandon and Michalakis 2002, Kawecki and Ebert 2004). For example, parasites are typically expected to be locally adapted to their hosts because of their shorter generation times, but low migration rates and low genetic diversity in the parasite can theoretically reverse this prediction (Greischar and Koskella 2007).

Parasites experience strong evolutionary pressures from both biotic and abiotic sources. Adapting to the physical environment may be equally as important as adapting to hosts, especially for parasites of ectotherms or those with life cycles involving free-living forms. Despite this, previous studies are limited in that they examine local adaptation to hosts or local adaptation to temperature, but not both. To our knowledge, only one prior study has demonstrated local adaptation of parasites to both hosts and temperature (Laine 2008), and was limited in its spatial extent with <1°C difference in temperature among populations. More typically, studies consider temperature as an ecological stressor rather than an evolutionary driver of local adaptation. In these cases, the evidence is mixed as to whether temperature significantly affects the pattern of host – parasite local adaptation (e.g., Blanford et al. 2002, Laine 2008) or whether there is a consistent pattern of local adaptation across temperature (e.g., Landis et al. 2012).

The western treehole mosquito *Aedes sierrensis* and its facultative ciliate parasite *Lambornella clarki* present an ideal system for investigating patterns of adaptation of a parasite to its host and to its thermal environment. *Ae. sierrensis* is a broadly distributed mosquito, inhabiting water-filled treeholes across the western United States. *Ae. sierrensis* is commonly infected by *L. clarki*, a facultative ciliate parasite that invades the host’s cuticle, replicates inside, and is released upon the death of its host. Both species occur across a broad latitudinal temperature gradient and can be easily reared in the lab. (Broberg and Bradshaw 1997, Hawley 1985, Washburn and Anderson 1986). Using this model system, we aim to test (i) whether parasitism is temperature dependent and (ii) whether parasites are locally adapted to hosts, temperature, or both. We employ a common garden experimental design that measures the interaction between parasite population, host population, and temperature in the laboratory.

## Materials and methods

### Collection of mosquito larvae and parasites

From November 2021 to April 2022, we collected larval *Ae. sierrensis* mosquitoes from field sites across California and Oregon to obtain individuals infected with *L. clarki*. For this experiment, we used populations of *Ae. sierrensis* and *L. clarki* collected from nine sites (Figure 1), which spans the majority of the range of *L. clarki* (Broberg and Bradshaw 1997, Washburn and Anderson 1986). The sites spanned from San Diego, CA to Portland, OR (33-45°N), and in elevation from coastal into the Sierra Nevada Mountains (1250 m elevation). The annual mean temperature ranged from 11-18°C and the average temperature of the wettest quarter—the activity period for *Ae. sierrensis* and *L. clarki* in treeholes—ranged from 5-13°C (coordinates and temperatures are given in Table S1). Once collected, mosquitoes were brought back to the lab, maintained in the dark at 7°C, and examined under a microscope for the presence of *L. clarki* infections. The parasite was then isolated by transferring infected mosquitoes into vials containing an autoclaved barley seed and 1 mL of autoclaved media made of 1 L distilled water, 1 protozoan pellet, and 0.38g of Herptivite (Fukami 2004). The *L. clarki* vials were kept in the dark at 21°C and new media was added weekly.

**Figure 1.**
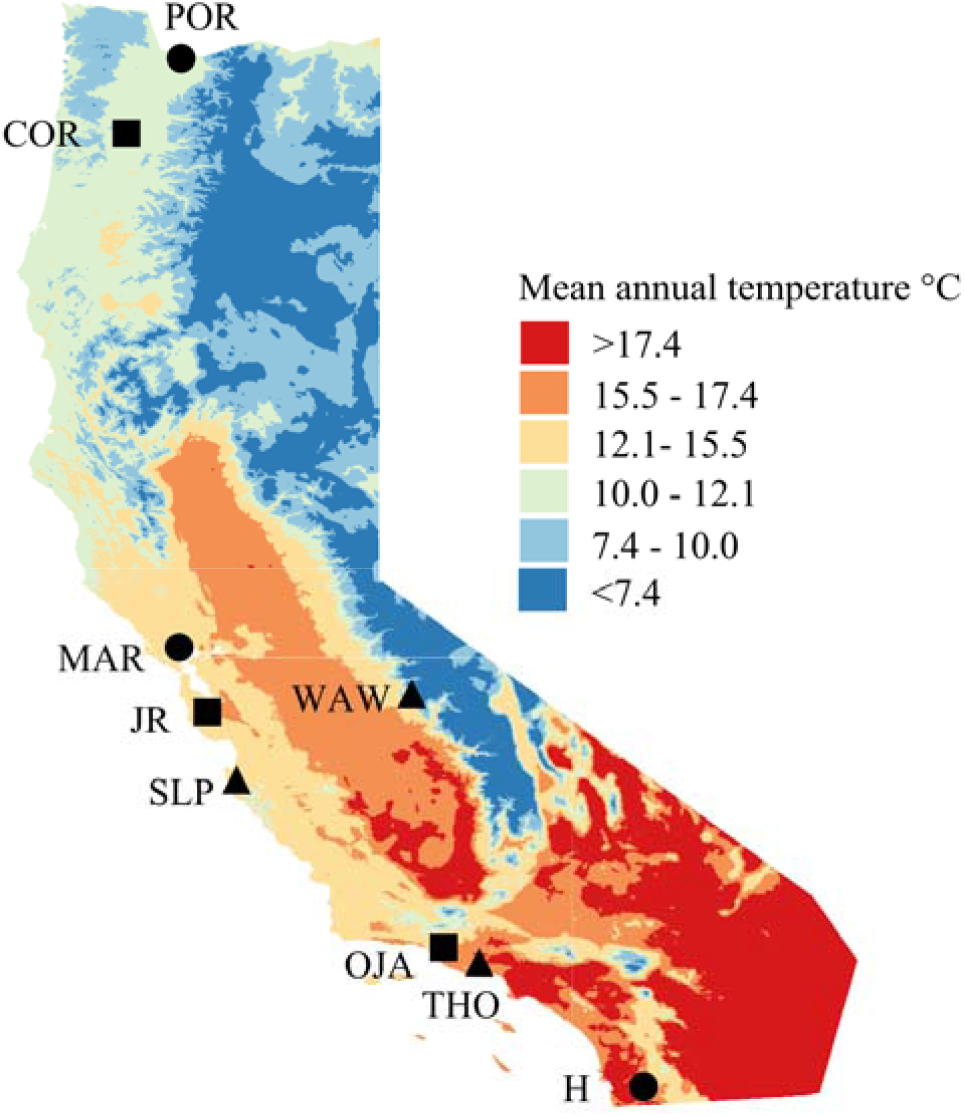
The nine populations used in the experiment span a wide climatic gradient and most of the species’ geographic range. The map is colored by mean annual temperature (°C). Point shapes represent the three groups used in the experiment, within which all possible host-parasite pairs were tested.

In order to obtain large numbers of first instar mosquito larvae used in the experiment, we reared field-collected mosquitoes to the next generation. Larval mosquitoes were brought back from the field and maintained in 100 mL plastic containers and fed a 4% solution of high-protein cat chow, 36% bovine liver powder, and 14% brewer’s yeast (Maïga 2017) weekly. We defined a population as mosquitoes and parasites from treeholes within a 20 km radius, which included between two and five treeholes, depending on the population. For each site we raised mosquitoes from all treeholes except for those where infected larvae were found. Once emergence began, adults were kept in collapsible aluminum cages (BioQuip, Rancho Dominguez, CA, USA) and fed a 10% sugar-water solution twice per week and defibrinated sheep’s blood once per week. Females laid eggs on damp filter paper placed inside black oviposition cups. Egg papers were labeled in plastic bags and maintained in the dark at 5°C for 1-3 months. To begin the experiment, eggs were hatched by submerging the egg papers in a solution of 500 mL Arrowhead distilled water, 300 mL autoclaved tree hole water, and 1 teaspoon Brewer’s yeast (Schwan and Anderson 1980).

### Infection experiment

We conducted a common garden experiment performed in the laboratory to examine patterns of parasite local adaptation to hosts and to temperature. We split the nine populations into three groups (Figure 1), each containing a warm site (annual mean temperature: 15.4-17.8°C, wettest quarter temperature: 11.3-13.2°C), a temperate site (annual 12.8-15.0°C, wettest quarter 9.6-10.5°C), and a cold site (annual 11.4-11.7°C, wettest quarter 5.3-5.6°C). To test for local adaptation of parasites to their hosts, each population of *L. clarki* was introduced into one sympatric and two allopatric populations of mosquito larvae. To test for local adaptation to temperature, we used three temperatures (7, 13, and 18°C), reflecting temperatures of the source sites. We had five replicates for each *L. clarki – Ae*.*sierrensis* pair by temperature treatment, except for OJA and POR, which had four replicates due to lower numbers of larvae that hatched.

We filled 6-well non-treated Falcon plates with five first-instar mosquito larvae in each well. To each well we introduced 4 mL of *L. clarki* at a standard inoculation concentration of 320 cells/mL. The plates were maintained in a 14:10 light-dark cycle and each well fed with 50 uL of 4% larval food suspension after each check for infection, which was chosen so that mosquitoes were not food limited (Maïga 2017). We monitored infection four times before the first day the larvae were expected to pupate. Because the development is faster at higher temperatures, the 18°C treatment was checked on days 3, 6, 9, and 12; the 13°C treatment was checked on days 5, 10, 15, and 20; and the 7°C treatment was checked on days 8, 16, 24, and 32.

During the examinations, we recorded the presence of cuticular cysts, melanization spots, internal infection, and survival. Cuticular cysts form as the parasite attacks the mosquito and melanization spots are the mosquito immune response to these cysts. As a final assay of infection, all dead larvae and larvae surviving through the fourth check were stained with black amide dye for 30 minutes as an additional check for *L. clarki* cysts on the cuticle of the the host (Soldo and Merlin 1972, Washburn et al. 1988). All mosquitoes were inspected under a dissecting microscope at 40X magnification.

### Free-living thermal performance experiment

As *L. clarki* is a facultative ciliate parasite that can complete its life cycle as a free-living organism, we also conducted an experiment to examine local adaptation of the free-living form to temperature. We measured the exponential growth rate of each population at seven temperatures: 5, 7, 12, 18, 21, 23, and 28°C. For each temperature and population of *L. clarki* we had five replicates, initiated by adding 2 mL of a low density culture (3 cell/100uL) and 1 barley seed to a 4 mL vial. We measured density daily for 6 days by plating 100 uL in small droplets on a petri dish and counting the number of cells.

### Statistical analysis

For the infection experiment, we report results of the number of larvae with cuticular cysts, melanization spots, and internal infection, as well as survival rates. To examine whether parasites are locally adapted to hosts, we included a binary independent variable for sympatric versus allopatric pairs. To examine whether parasites are locally adapted to temperature, we included a binary independent variable for a parasite in the experimental condition that was matched to its source environment versus one that was mismatched. A mismatched experimental condition was one in which a population was exposed to a temperature typically outside of its normal range based on mean annual and wettest quarter temperatures (e.g., a cold population is matched to 7°C and mismatched to 13 and 18°C). To examine if there was an effect of parasite population, host population, or temperature, these terms were also included as categorical variables in the model.

We ran generalized linear models using the glm function in the “stats” package in R version 4.1.1. We ran Poisson regressions for the response variables: number of larvae with cuticular cysts, number of larvae with melanization spots, and number of larvae with internal infection. Because these responses happen sequentially, we chose to analyze the time point when each response peaked (i.e., checkpoint 1 for cysts, checkpoint 2 for melanization, and checkpoint 4 for internal infection). For survival, we calculated the percentage of all larvae that had died by checkpoint 4 and because this percentage was so low we did not conduct further analysis on this variable. For each response variable, our main model included parasite population, host population, temperature, sympatry/allopatry, and matching/mismatching temperature.

Additionally, to examine whether there was a temperature-mediated effect on local adaptation to hosts, we also included an interaction between temperature and sympatry/allopatry, but this did not significantly improve the model and it was dropped. We determined significance using likelihood ratio tests and report deviance and p-values.

For the growth rate experiment, we subsetted the data to only include days one through five because by day six, the rate of growth at many temperatures was slowing down. We then calculated the exponential growth rate for each replicate using “nls” in R. We fit thermal performance curves to these growth rates using the Rezende 2019 model as this model visually produced a good fit to all the populations and bootstrapped to get 95% confidence intervals using the “rTPC” package (Rezende and Bozinovic 2019, Padfield et al. 2021). To test for local adaptation to temperature we calculated Pearson’s correlation coefficient between annual mean temperature of the site (Table S1) and peak growth rate temperature.

## Results

### Documenting the infection pathway

We documented the pathway of infection, in which parasites were introduced as free-living *L. clarki*, those cells rapidly transformed to attack the cuticle of the mosquito larvae, the mosquitoes responded with a melanization response, larvae were infected internally, and larvae died of infection (Figure 2). As expected, the proportion of wells containing free-living *L. clarki* declined over time. By the first checkpoint, infective cysts were present and we observed the highest number of larvae with cuticular cysts. At the second checkpoint we observed the highest number of larvae with melanization spots. The number of infected larvae steadily rose over time. These temporal trends were the same across temperature treatments, except for cuticular cysts, which peaked at the second checkpoint for the 7°C and 18°C treatments, and for melanization, which peaked at the third checkpoint for the 7°C treatment (Figure S1). Because so few larvae died by the end of the experiment (11%), we did not use larval survival as a response variable in any further analysis.

**Fig 2.**
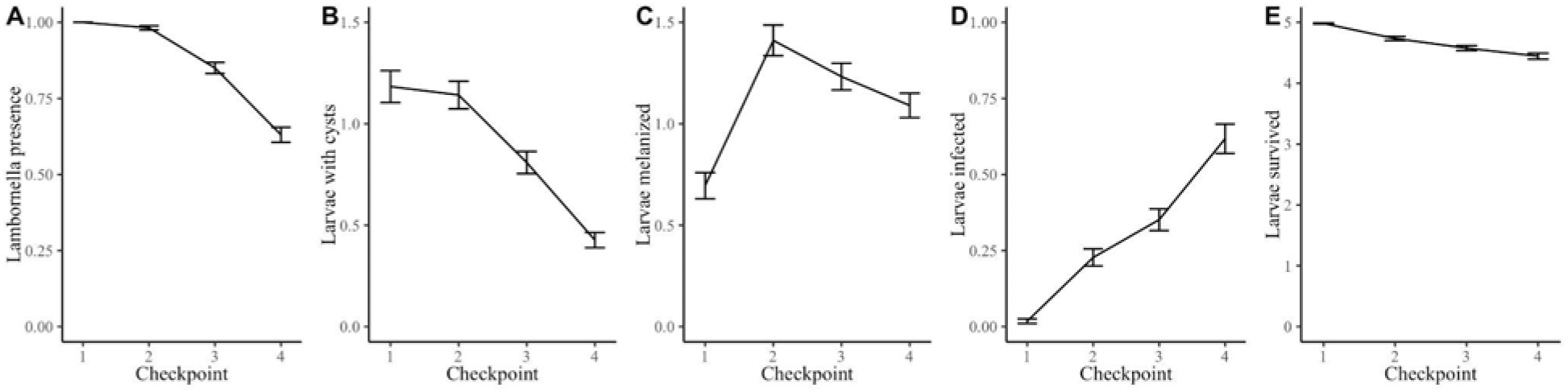
The pathway of infection over time. *L. clarki* presence (A) and cysts (B) peaked at the first checkpoint. Melanization (C) peaked at the second checkpoint. Infection (D) increased over time and survival (E) decreased over time. Lines represent averages across all population pairs, temperatures, and replicates; error bars represent ±1 SE.

### Local adaptation

We found that parasites are locally adapted to their hosts (Figure 3, see Figure S2 for individual parasite populations). Sympatric populations had 3.2 times more larvae with cysts (□^2^ (1)=15, p<0.001) and 2.6 times more larvae that were infected (^2^ (1)=62, P<0.001) compared to allopatric populations. Also, sympatric populations had fewer larvae that were melanized (□^2^(1)=3.8, P=0.05). Hence, not only did *L. clarki* infect sympatric hosts at a higher rate than allopatric hosts, but also sympatric *L. clarki* elicited a weaker immune response from the host. We also examined the relationship between the number of larvae infected and the geographic distance and the climate distance between host and parasite pairs, because we expect that as pairs become more distant or climatically dissimilar, infection should decline. As expected, the number of infected larvae decreased with both climatic distance (R^2^= 0.12, P=0.04) and geographic distance between host and parasite, although the latter was not statistically significant (R^2^= 0.09, P=0.08). Additionally, there was no evidence that temperature mediates the signal of local adaptation to hosts: the number of larvae infected in sympatric populations was higher than the number infected in allopatric populations at all temperatures (Figure S4).

**Fig 3.**
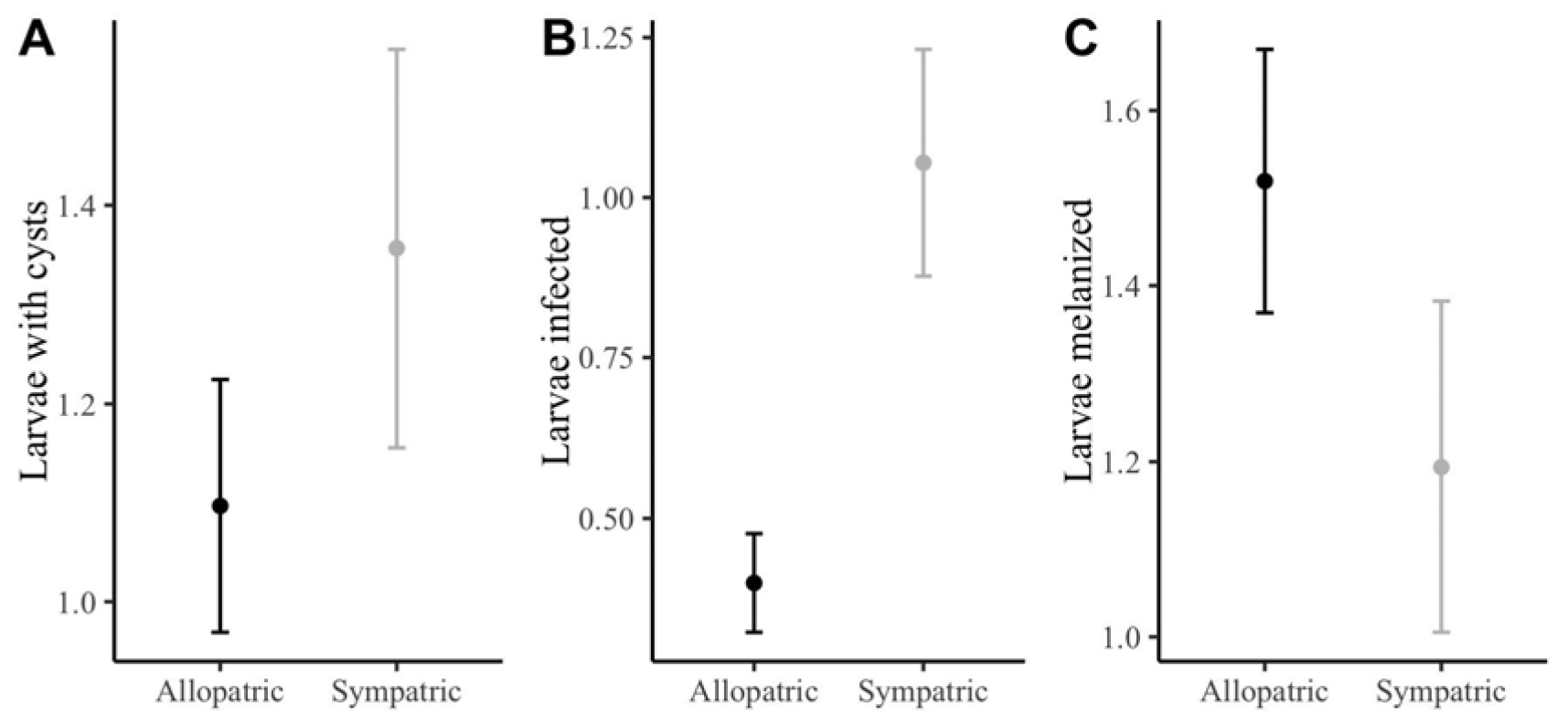
Parasites are locally adapted to hosts. The number of larvae with parasitic cysts (A) and infection (B) was higher in sympatric than allopatric populations. The number of larvae melanized (C), a sign of immune response, was higher in allopatric than sympatric populations. Points represent averages across all replicates and temperature treatments; error bars represent ±1 SE.

Including an interaction term between temperature condition in the experiment and the binary sympatry variable did not significantly improve the model (□^2^(2)=2.3, P=0.31).

In contrast to the signal of host adaptation, parasites were not locally adapted to temperature (Figure 4, see Figure S5 for individual parasite populations). There were no significant differences in the number of larvae with cysts (□^2^(1)=0.12, P=0.73), the number of larvae that were infected (□^2^(1)=0.74, P=0.39), or the number of larvae melanized (□^2^(1)=0.23, P=0.63) between parasites that were placed at temperature that matched their source environment versus those placed at mismatched temperatures (Fig. 4).

**Fig 4.**
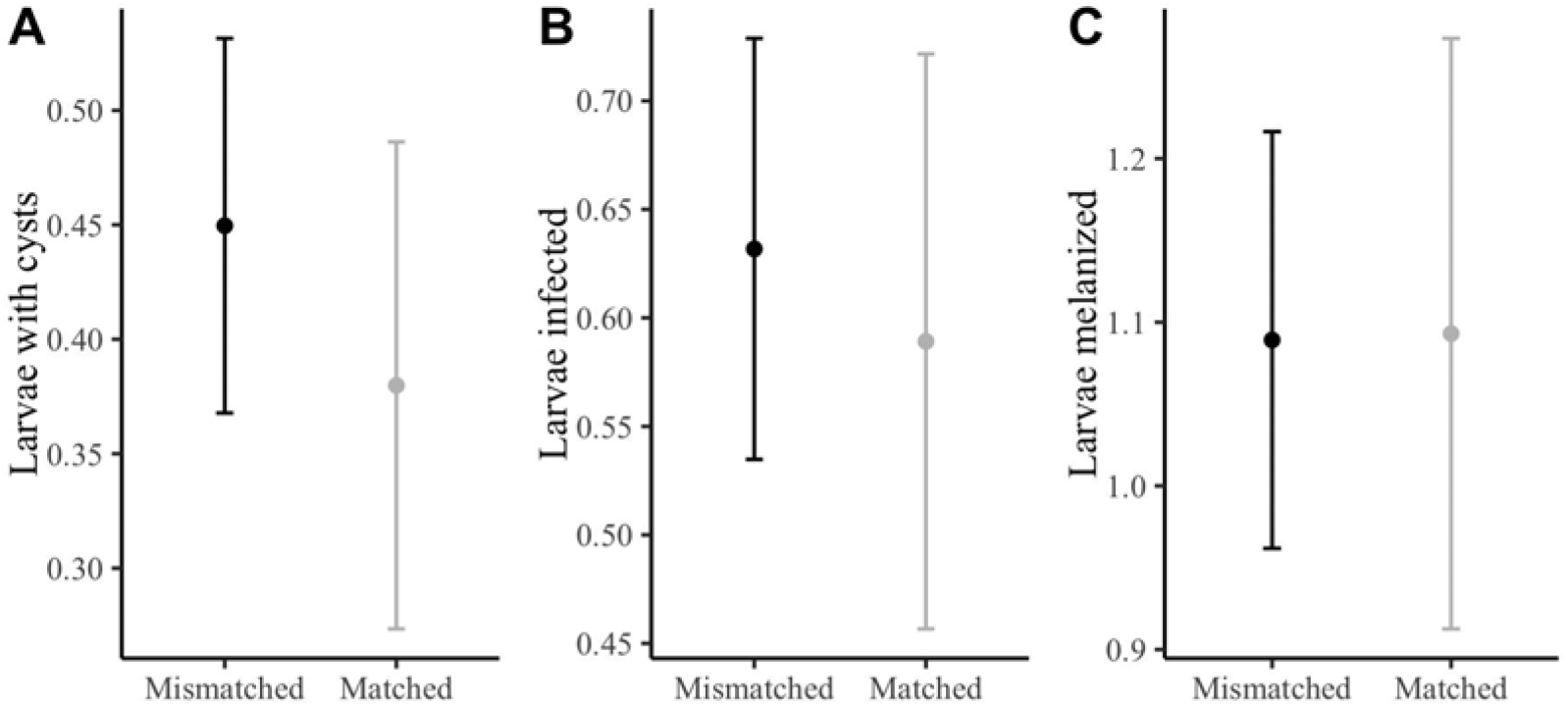
Parasitic *L. clarki* are not locally adapted to temperature. The number of larvae with cysts (A), infection (B), and melanization (C) did not significantly differ between populations that were matched versus mismatched to their source environment. Points represent averages across all replicates and sympatric/allopatric population pairs; error bars represent ±1 SE.

Parasite populations varied significantly in infection success (Figure 5A, Figure S6). This is often referred to as a ‘deme effect’ in the local adaptation literature, signifying that certain populations are overall more fit than others, regardless of habitat. Parasites varied in cysts □^2^ (6)=344, P<0.001), melanization (□^2^ (6)=53, P<0.001), and infection (□^2^ (6)=269, P<0.001).

**Figure 5.**
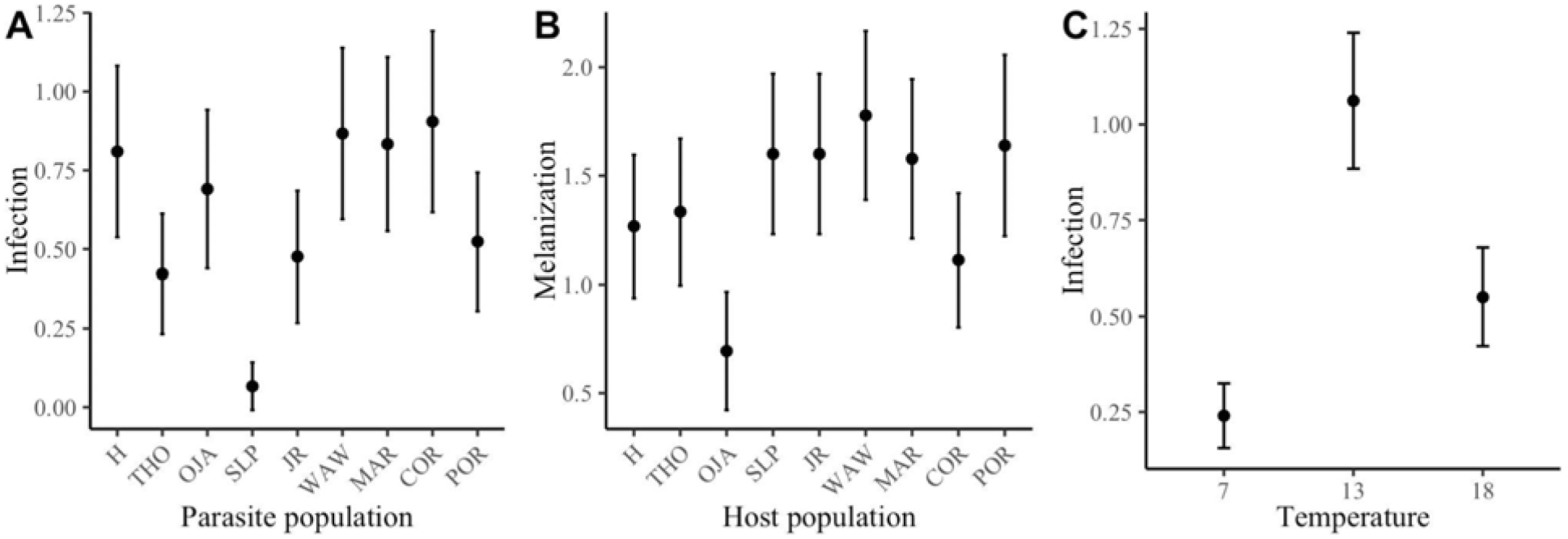
There is significant variation in deme and habitat quality. Variation in A) infection rates between parasite populations, averaged across temperatures, host populations, and replicates, B) melanization between host populations, averaged across parasite populations, temperatures, and replicates, and C) infection rates across temperature, averaged across host and parasite populations and replicates. Error bars represent ±1 SE.

We found that one population (‘SLP’) performed especially poorly, producing the lowest number of infections of all nine populations. Host populations also varied significantly in cysts □^2^ (6)=22, P<0.001; Figure S6), melanization (□ ^2^(6)=18, P<0.01; Figure 5B), and infection □^2^ (6)=36, P<0.001; Figure S6). Host population can be considered one aspect of habitat quality, with certain host populations being more or less resistant to infection. Although there was variation in both host and parasite populations, this did not correlate with the latitudinal or temperature gradient from which they were collected.

Temperature significantly affected infection, with peak infection occurring at the intermediate temperature of 13°C (□^2^(2)=74, P<.001; Figure 5C), as expected from principles of thermal biology. Temperature also significantly affected cysts and melanization, with the highest number of larvae with cysts occurring at 13°C and the highest number of larvae melanized occurring at the warmest temperature of 18°C (□^2^(2)=15, P<.001; Figure S6).

Although there was no indication that parasitic *L. clarki* are locally adapted to temperature, we did find evidence that free-living *L. clarki* may be locally adapted to temperature. Growth rates of free-living *L. clarki* from warmer sites peaked at higher temperatures than those from colder temperatures (Figure 6, Figure S7). Peak growth rate temperature was strongly positively correlated with annual mean temperature of the collection site (r = 0.80). Moreover, the temperatures of peak free-living ciliate growth (16.5-23.5°C) were substantially higher than the temperature that maximized infection in the experiment (13°C).

**Figure 6.**
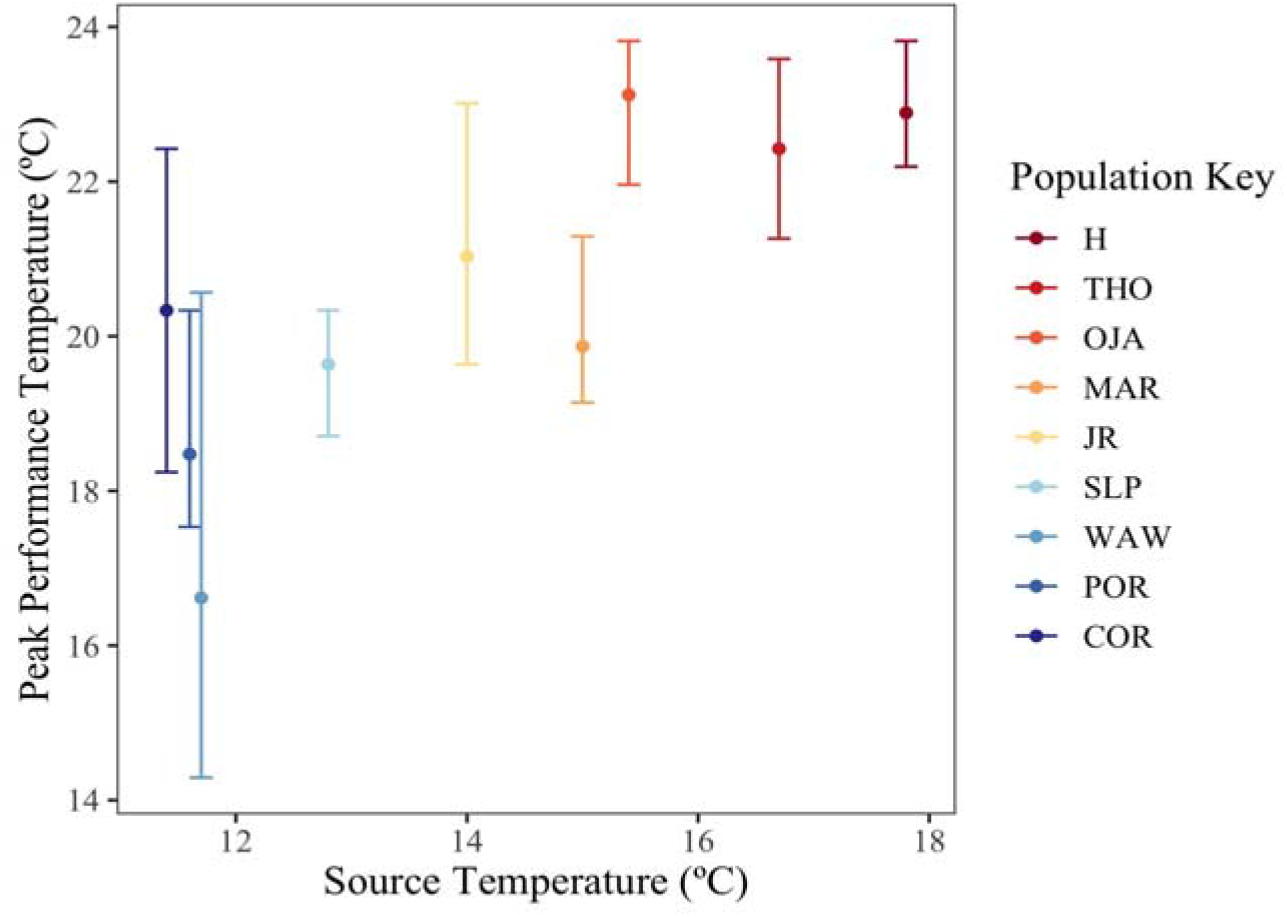
Free-living forms of *L. clarki* show a signal of local adaptation to temperature, in contrast to parasitic forms. The peak growth rate temperature (y-axis) is positively correlated (r = 0.80) with the mean annual temperature of the parasite’s source environment (x-axis). Populations are colored in order of decreasing mean annual temperature. Error bars represent bootstrapped 95% confidence intervals.

## Discussion

Our study examined the extent to which parasites are adapted to their biotic and abiotic environments, revealing that parasites are locally adapted to their hosts but not to temperature (Figures 3 and 4). This supports the theoretical expectation that most parasites are ahead in the evolutionary arms race due to their shorter generation times than hosts. At the same time, our finding that *L. clarki* is adapted to its mosquito host is surprising for several reasons. First, a review of experiments found that parasite local adaptation is relatively rare, only seen in roughly a third of cases (Greischar and Koskella 2007). In contrast to many previous studies analyzed in the review, our experimental design includes many populations (N = 9), spanned the entire range of the focal organism, and tested the interaction under multiple environmental conditions, providing a more comprehensive picture of parasite local adaptation. Second, *L. clarki* is a facultative parasite. Therefore, despite existing and reproducing as a free-living organism in the absence of mosquitoes, it experiences such strong selective pressure from mosquito predation that it has evolved a parasitic stage that has further adapted locally to mosquito populations.

Third, given that local adaptation is influenced by the relative migration rates of host and parasite, it is noteworthy that *L. clarki* parasites are likely more dispersal limited than their host, which has a flying adult life stage. Parasites are transported to another treehole on rare occasions in which they infect the reproductive tract of an adult female mosquito (Egerter and Anderson 1985), while dispersal strategies of free-living *L. clarki* are unknown. Despite greater host dispersal, the parasite is ahead in the evolutionary arms race.

The pattern of parasites being locally adapted to hosts was consistent across temperatures. That is, parasites caused higher infection in sympatric, relative to allopatric hosts, across all temperature treatments (Figure S4). This result aligns with previous findings in a fish – trematode parasite system, in which the parasite is evolutionarily ahead of the host even under heat stress (Landis et al. 2012). However, in other systems such as a pea aphid fungal pathogen (Blanford et al. 2002) and a plant fungal pathogen (Laine 2008), temperature mediated patterns of local adaptation, even changing the direction of adaptation. *L. clarki* parasites appear to be adapting to hosts both by increasing their attack rates, evidenced by more sympatric larvae with cysts, and by evading the immune response of the host, evidenced by fewer sympatric larvae with melanization.

Despite inhabiting locations that vary by over 8°C in their annual mean temperature, we did not find any evidence that *L. clarki* populations are locally adapted to temperature in their parasitic form (Figure 4). This suggests that *L. clarki* parasites may face stronger selective pressures from hosts than from the thermal environment. In contrast, we found a signal of local adaptation to temperature in free-living *L. clarki* growth rates (Figure 6), suggesting that temperature is an important selective pressure during free-living phases of the ciliate life cycle. However, one limitation of our study was the small number of temperature treatments. By only testing three temperatures, we may have missed fine scale differences in local adaptation of infection to temperature, e.g., if the optimum temperatures for infection varied by only a few degrees between populations (but note that the optimal temperatures for free-living growth varies by ∼7°C among populations).

When we consider the performance of parasite populations on average across treatments, we find that some populations have consistently high attack and infection rates while others have consistently low attack and infection rates (Figure 5) (i.e., a ‘deme quality effect’, Blanquart et al. 2013). Populations that perform poorly in any environment may have poor genetic quality due to factors such as inbreeding and drift in small populations, or they may be poorly adapted to the laboratory experimental environment. Although we may expect populations near the range edge, such as our Northernmost (POR) or Southernmost (H) population to perform poorly relative to those in the center (Bontrager et al. 2021), we did not find evidence of this nor any consistent pattern in deme quality across space. We also considered that a low fitness parasite population might be better adapted to free-living growth than to infection, but we did not detect a tradeoff between infection success and maximum free-living growth rate. In addition to variation among parasite populations, we also find variation among host populations in their susceptibility to attack and infection and in their melanization response (Figure 5). Similar to parasites, we did not find a consistent pattern in host quality across space. Host – parasite and predator – prey systems can also display geographic mosaics of hot spots and cold spots of coevolution rather than a continuous gradient (Thompson 2005). For example, in the classic case study of newts and garter snakes, areas where newts are the most toxic coincide with areas where garter snakes are most resistant to the toxin (Brodie et al. 2002). However, in this study we find that locations with the most resistant hosts are not the locations with the most infective parasites. Going forward, to fully understand the coevolutionary dynamics between host and parasite, obtaining multiple years of data or performing a time-shift experiment in which hosts are exposed to parasites from a different time point, will be necessary (Penczykowski et al. 2016, Thompson 1999).

Overall, we find that infection is temperature-dependent, peaking at the intermediate temperature of 13°C. For most ectotherms, performance declines away from the thermal optimum towards their thermal limits, and parasites are no exception (Lafferty and Mordecai 2016). Although our experiment is limited to three temperature treatments, we capture a nonlinear relationship between temperature and infection for *L. clarki*. We observed a peak infection temperature that was lower than the peak temperature for free-living *L. clarki* growth rates (i.e., 13°C vs 16.5-23.5°C). Because infection involves the thermal performance of both parasite and host (Gehman et al. 2018), the difference in optimal temperatures between free-living growth and parasite infection may be explained by the relative influence of temperature on parasite attack versus host defense. Specifically, the melanization response of mosquitoes increased with temperature, which adds support to other studies demonstrating the temperature dependence of insect immunity (Murdock et al. 2012). A consequence of a hump-shaped relationship with temperature is that we should not expect climate change to consistently increase or decrease disease prevalence (Rohr et al. 2011). Instead, the effects of warming on infection will differ across a species range, with climate change potentially driving geographic shifts in prevalence such as that observed in white pine blister rust (Dudney et al. 2021).

In summary, our study documents a coevolutionary interaction between a ciliate parasite and its mosquito host in the context of a temperature gradient, where the parasite is ahead in the evolutionary arms race. Despite a large body of literature on host–parasite coevolutionary arms races and theory suggesting that these interactions drive evolutionary and population dynamics in both players, our experiment provides some of the first clear evidence of rangewide parasite local adaptation to its host. Although the parasite is ahead in the arms race, the host also plays a role in the outcome of the interaction, evidenced by a much lower optimal temperature for parasite infection than for free-living ciliate growth. The host immune response, which peaks at higher temperatures, appears to moderate the high free-living growth rates at warm temperatures and drive down the thermal optimum for parasitism. Interestingly, we did not find evidence that *L. clarki* is locally adapted to temperature in its parasitic form, despite finding this evidence for its free-living form. Adaptation of *L. clarki* parasites to hosts but not temperature suggests that the selective pressure from hosts may be stronger than that of the thermal environment. We also captured a nonlinear response of infection to temperature, demonstrating that although there was a lack of local thermal adaptation, temperature remains an important factor mediating host– parasite interactions.

## Acknowledgements

This work was supported by the NSF Postdoctoral Research Fellowships in Biology Program under Grant No. 2208947. EAM was supported by the National Institutes of Health (R35GM133439, R01AI168097, R01AI102918), the National Science Foundation (DEB-2011147, with Fogarty International Center), and the Stanford Center for Innovation in Global Health, King Center on Global Development, and Woods Institute for the Environment. LIC was funded by the Philippe Cohen Graduate Fellowship and the Stanford Center for Computational, Evolutionary and Human Genomics. JEF was funded by the Bing-Mooney Fellowship. We would like to thank the many agencies in California, including Placer, San Diego, San Mateo, Santa Barbara, and Riverside mosquito and vector control districts, who assisted us in locating mosquito populations and specifically, Bret Barner and Angie Nakano for sharing animal husbandry information. We would also like to thank members of the Mordecai lab who helped set up the experiment.

## References

Altizer, S., Ostfeld, R. S., Johnson, P. T., Kutz, S., & Harvell, C. D. (2013). Climate change and infectious diseases: from evidence to a predictive framework. Science, 341(6145), 514-519.

Blanford, S., Thomas, M. B., Pugh, C., & Pell, J. K. (2003). Temperature checks the Red Queen? Resistance and virulence in a fluctuating environment. Ecology Letters, 6(1), 2–5.

Blanquart, F., Kaltz, O., Nuismer, S. L., & Gandon, S. (2013). A practical guide to measuring local adaptation. Ecology Letters, 16(9), 1195–1205.

Bontrager, M., Usui, T., Lee□Yaw, J. A., Anstett, D. N., Branch, H. A., Hargreaves, A. L., et al. (2021). Adaptation across geographic ranges is consistent with strong selection in marginal climates and legacies of range expansion. Evolution, 75(6),1316–1333.

Broberg, L. E., & Bradshaw, W. E. (1997). Lambornella clarki (Ciliophora: Tetrahymenidae) resistance in Aedes sierrensis (Diptera: Culicidae): population differentiation in a quantitative trait and allozyme loci. Journal of Medical Entomology, 34(1), 38-45.

Brodie Jr, E. D., Ridenhour, B. J., & III Brodie, E. D. (2002). The evolutionary response of predators to dangerous prey: hotspots and coldspots in the geographic mosaic of coevolution between garter snakes and newts. Evolution, 56(10), 2067–2082.

Dudney, J., Willing, C. E., Das, A. J., Latimer, A. M., Nesmith, J. C., & Battles, J. J. (2021). Nonlinear shifts in infectious rust disease due to climate change. Nature Communications, 12(1),5102.

Egerter, D. E., & Anderson, J. R. (1985). Infection of the western treehole mosquito, Aedes sierrensis (Diptera: Culicidae), with Lambornella clarki (Ciliophora: Tetrahymenidae). Journal of Invertebrate Pathology, 46(3), 296–304.

Fukami, T. (2004). Assembly history interacts with ecosystem size to influence species diversity. Ecology, 85(12),3234–3242.

Gandon, S. (2002). Local adaptation and the geometry of host–parasite coevolution. Ecology Letters, 5(2), 246–256.

Gandon, S., & Michalakis,Y. (2002). Local adaptation, evolutionary potential and host–parasite coevolution: interactions between migration, mutation, population size and generation time. Journal of Evolutionary Biology, 15, 451–462.

Gehman, A. L. M., Hall, R. J., & Byers, J. E. (2018). Host and parasite thermal ecology jointly determine the effect of climate warming on epidemic dynamics. Proceedings of the National Academy of Sciences, 115(4),744–749.

Greischar, M. A., Koskella, B. (2007). A synthesis of experimental work on parasite local adaptation. Ecology Letters, 10(5), 8–434.

Hawley, W. A. (1985). The effect of larval density on adult longevity of a mosquito, Aedes sierrensis: epidemiological consequences. The Journal of Animal Ecology, 955–964.

Kawecki, T. J., & Ebert, D. (2004). Conceptual issues in local adaptation. Ecology Letters, 7(12), 1225–1241.

Laine, A. L. (2008). Temperature-mediated patterns of local adaptation in a natural plant– pathogen metapopulation. Ecology Letters, 11(4),327–337.

Lafferty, K. D., & Mordecai, E. A. (2016). The rise and fall of infectious disease in a warmer world. F1000Research, 5.

Landis, S. H., Kalbe, M., Reusch, T. B., & Roth, O. (2012). Consistent pattern of local adaptation during an experimental heat wave in a pipefish-trematode host-parasite system. PLoS One, 7(1),p e30658.

Maïga, H. (2017). Guidelines for Routine Colony Maintenance of Aedes Mosquito Species: FAO-IAEA.

Murdock, C. C., Paaijmans, K. P., Bell, A. S., King, J. G., Hillyer, J. F., Read, A. F., & Thomas, M. B. (2012). Complex effects of temperature on mosquito immune function. Proceedings of the Royal Society B: Biological Sciences, 279(1741), 3357–3366.

Padfield, D., O’Sullivan, H., & Pawar, S. (2021). rTPC and nls.multstart: A new pipeline to fit thermal performance curves in R. Methods in Ecology and Evolution. https://doi.org/10.1111/2041-210X.13585

Penczykowski, R. M., Laine, A. L., & Koskella, B. (2016). Understanding the ecology and evolution of host–parasite interactions across scales. Evolutionary Applications, 9(1), 37–52.

Rezende, E. L., & Bozinovic, F. (2019). Thermal performance across levels of biological organization. Philosophical Transactions of the Royal Society B, 374(1778), 20180549.

Rohr, J. R., Dobson, A. P., Johnson, P. T., Kilpatrick, A. M., Paull, S. H., Raffel, T. R., et al. (2011). Frontiers in climate change–disease research. Trends in Ecology & Evolution, 26(6), 270–277.

Schwan T.G., & Anderson, A.R. (1980). A comparison of egg hatching techniques for the western treehole mosquito, Aedes sierrensis. Mosquito News 0:263–269.

Stevenson, L.A., Alford, R.A., Bell, S.C., Roznik, E.A., Berger, L. & Pike, D.A. (2013). Variation in thermal performance of a widespread pathogen, the amphibian chytrid fungus Batrachochytrium dendrobatidis. PloS one, 8(9),p p.e73830.

Sternberg, E. D., & Thomas, M. B. (2014). Local adaptation to temperature and the implications for vector-borne diseases. Trends in Parasitology 30:115–122.

Thompson, J. N. (1999). Specific hypotheses on the geographic mosaic of coevolution. The American Naturalist, 153(S5), S1–S14.

Thompson, J. N. (2005). The geographic mosaic of coevolution. In The geographic mosaic of coevolution. University of Chicago Press.

Washburn, J. O., & Anderson, J. R. (1986). Distribution of Lambornella clarki (Ciliophora: Tetrahymenidae) and other mosquito parasites in California treeholes. Journal of Invertebrate Pathology, 48(3), 296-309.

Washburn, J. O., Gross, M. E., Mercer, D. R., & Anderson, J. R. (1988). Predator-induced trophic shift of a free-living ciliate: parasitism of mosquito larvae by their prey. Science, 240(4856), 1193–1195.

Soldo, A. T., & Merlin, E. J. (1972). The cultivation of symbiote□free marine ciliates in axenic medium. The Journal of Protozoology, 19(3), 519–524.

