## Supplement for "A mosquito parasite is locally adapted to its host but not temperature"

Supplemental Information for "A mosquito parasite is locally adapted to its host but not temperature"

**Table S1.** Climate variables for the 9 populations. For each population, coordinates are given for the two to five treeholes from which mosquitoes were collected, along with the centroid of the points. Mean annual temperature and mean temperature of the wettest quarter in the bioclim dataset (WORLDCLIM; Fick and Hijmans 2017) were extracted for these coordinates and a mean is provided for each population.

| **Treehole** | **Latitude** | **Longitude** | **Mean annual temperature °C** | **Mean temperature wettest quarter °C** |
|---|---|---|---|---|
| H01 | 32.6800003 | -116.82 | 17.8 | 13.2 |
| H02 | 32.6899986 | -116.82 | 17.7 | 13.1 |
| H04 | 32.6800003 | -116.82 | 17.8 | 13.2 |
| H06 | 32.6800003 | -116.82 | 17.8 | 13.2 |
| H08 | 32.6800003 | -116.82 | 17.8 | 13.2 |
| H07* | 32.6819229 | -116.81861 | 17.8 | 13.2 |
| **H centroid** | **32.6819871** | **-116.81977** | **17.8** | **13.2** |
| OJA03 | 34.5200005 | -119.27 | 14.7 | 9.9 |
| OJA05 | 34.416122 | -119.163033 | 15.0 | 10.3 |
| OJA06 | 34.3600006 | -119.32 | 15.9 | 12.5 |
| OJA07* | 34.3567726 | -119.31553 | 15.9 | 12.5 |
| **OJA centroid** | **34.41322365** | **-119.26713** | **15.4** | **11.3** |
| THO04 | 34.1699982 | -118.88 | 16.6 | 12.6 |
| THO06 | 34.2099991 | -118.82 | 16.8 | 12.5 |
| THO02* | 34.1679055 | -118.86089 | 16.7 | 12.7 |
| **THO centroid** | **34.1826343** | **-118.85363** | **16.7** | **12.6** |
| MAR26 | 38.1500015 | -122.58 | 14.8 | 9.6 |

| | | | | |
|---|---|---|---|---|
| MAR29 | 38.1500015 | -122.57 | 15.0 | 9.7 |
| MAR32 | 38.1300011 | -122.54 | 15.1 | 9.7 |
| MAR33 | 38.1300011 | -122.54 | 15.1 | 9.7 |
| MAR36 | 38.1300011 | -122.54 | 15.1 | 9.7 |
| MAR35* | 38.1304913 | -122.54399 | 15.1 | 9.7 |
| **MAR centroid** | **38.1367496** | **-122.55233** | **15.0** | **9.7** |
| JR04 | 37.4099998 | -122.23 | 14.0 | 10.5 |
| JR20 | 37.4000015 | -122.24 | 13.9 | 10.4 |
| JR05* | 37.4091187 | -122.23321 | 14.1 | 10.5 |
| **JR centroid** | **37.4063733** | **-122.2344** | **14.0** | **10.5** |
| SLP01 | 36.4500008 | -121.81 | 12.8 | 9.4 |
| SLP06 | 36.4700012 | -121.82 | 12.7 | 9.4 |
| SLP08* | 36.467147 | -121.83617 | 12.9 | 9.8 |
| **SLP centroid** | **36.462383** | **-121.82205** | **12.8** | **9.6** |
| POR02 | 45.5900002 | -122.38 | 11.6 | 5.5 |
| POR03 | 45.6800003 | -122.74 | 11.4 | 5.5 |
| POR07 | 45.5400009 | -122.65 | 11.8 | 5.4 |
| POR10 | 45.4900017 | -122.5 | 11.6 | 5.5 |
| POR09* | 45.446151 | -122.5076 | 11.3 | 5.3 |
| **POR centroid** | **45.5492308** | **-122.55552** | **11.6** | **5.4** |
| COR01 | 44.5400009 | -123.25 | 11.5 | 5.6 |
| COR02 | 44.5499992 | -123.27 | 11.6 | 5.6 |
| COR04 | 44.5600014 | -123.27 | 11.6 | 5.6 |
| COR09 | 44.6100006 | -123.28 | 11.2 | 5.4 |

| | | | | |
|---|---|---|---|---|
| COR08* | 44.6064994 | -123.27997 | 11.2 | 5.4 |
| **COR centroid** | **44.5733003** | **-123.26999** | **11.4** | **5.6** |
| WAW02 | 37.5499992 | -119.68 | 11.9 | 5.6 |
| WAW04* | 37.5426051 | -119.62716 | 11.4 | 5 |
| **WAW centroid** | **37.5463022** | **-119.65358** | **11.7** | **5.3** |

* Locations where *L. clarki* were collected

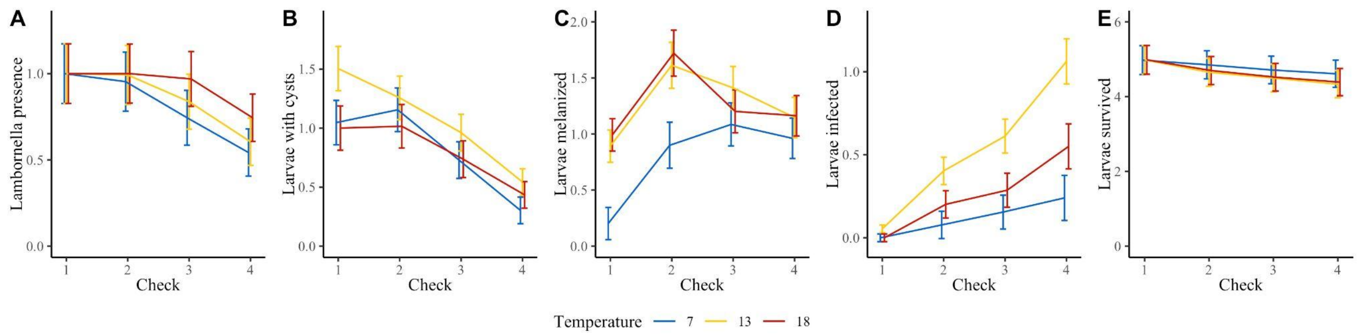


Figure S1. *L. clarki* presence (A), cysts (B), melanization (C), infection (D), and survival (E) over time, colored by temperature.

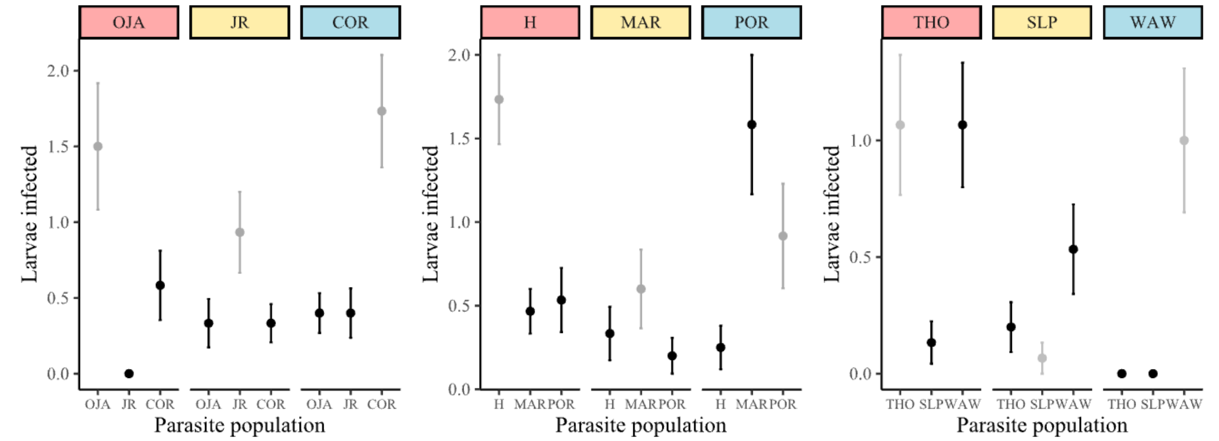


Figure S2. Infection rates of each pair of hosts and parasites, averaged over temperature. Host population is labeled above, colored as a warm (red), temperate (yellow), or cold (blue) site. Points are colored as sympatric (gray) or allopatric (black) pairs.

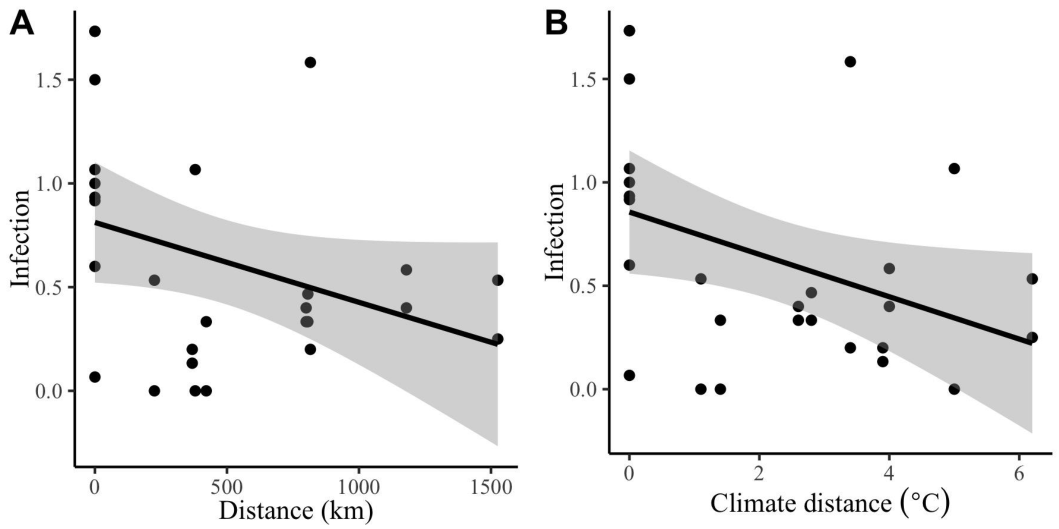


Figure S3. Infection decreases with (A) increasing geographic distance and (B) increasing climate distance between host–parasite pairs. The relationship with climate distance is significant and the relationship with geographic distance is not. Regression lines are in black with 95% confidence intervals in gray.

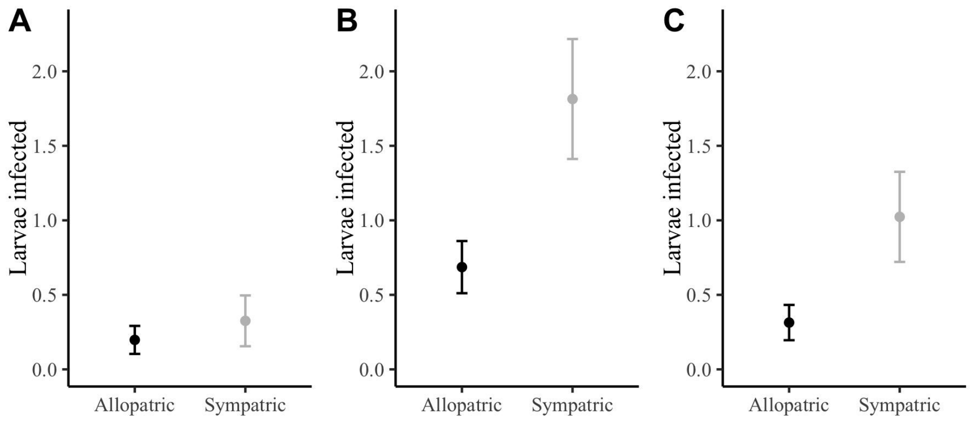


Figure S4. A comparison of infection in allopatric and sympatric populations at 7°C (A), 13°C (B), and 18°C (C). The interaction between temperature and local adaptation to hosts was not significant.

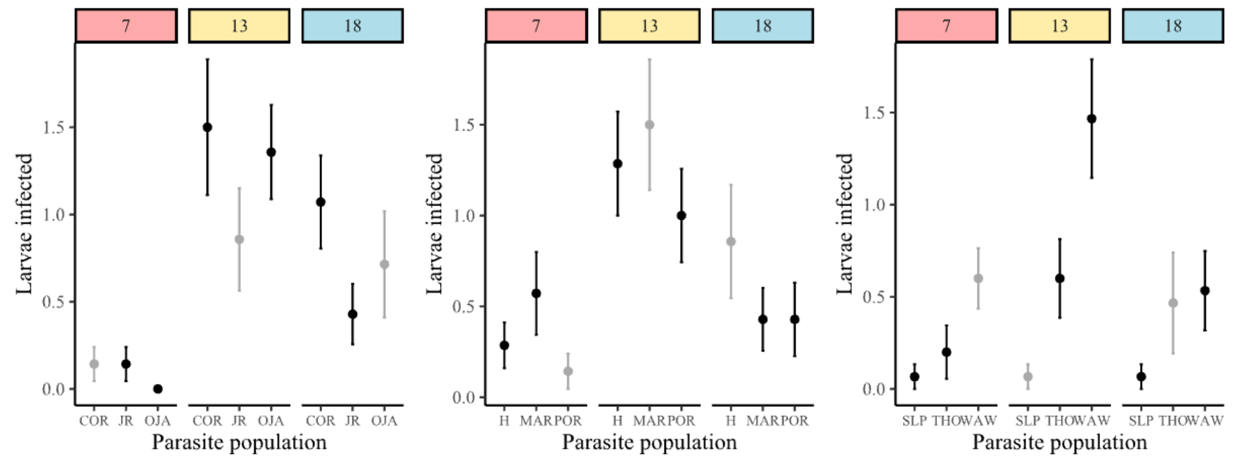


Figure S5. Infection rates of parasite populations plotted against temperature, averaged over host populations. The temperature of the experiment is labeled above as 7°C (red), 13°C (yellow), or 18°C (blue). Points are colored as sympatric (gray) or allopatric (black) pairs.

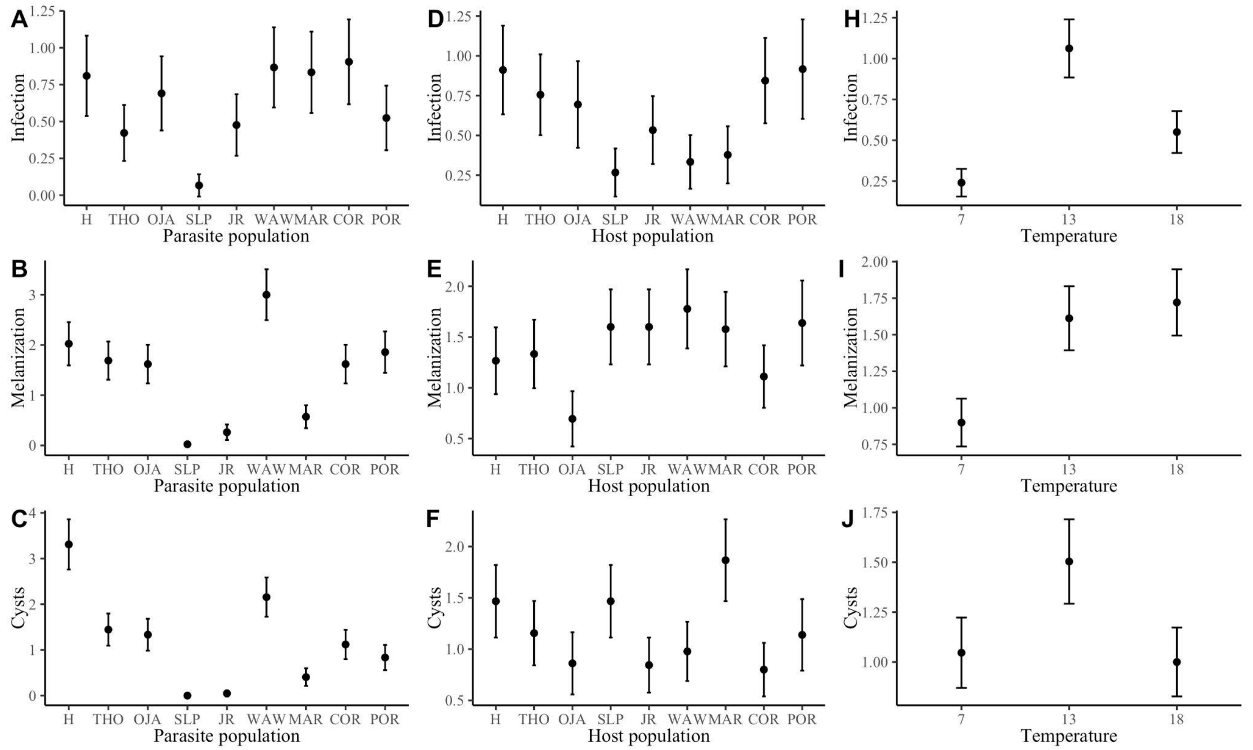


Figure S6. Variation in cysts, melanization, and infection among parasite populations (A-C), host populations (D-F), and temperature (H-J). Populations are ordered by increasing latitude.

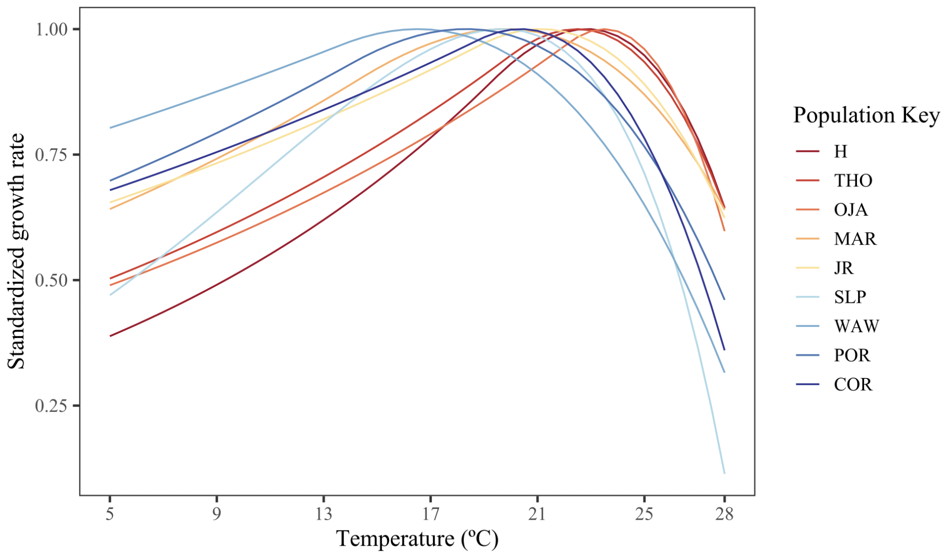


Figure S7. Thermal performance curves of the free-living *L. clarki*, with populations colored in order of decreasing mean annual temperature.
